# Right Ventricular Sarcomere Contractile Depression and the Role of Thick Filament Activation in Human Heart Failure with Pulmonary Hypertension

**DOI:** 10.1101/2023.03.09.531988

**Authors:** Vivek Jani, M. Imran Aslam, Axel J. Fenwick, Weikang Ma, Henry Gong, Gregory Milburn, Devin Nissen, Ilton Cubero Salazar, Olivia Hanselman, Monica Mukherjee, Marc K. Halushka, Kenneth B. Margulies, Kenneth Campbell, Thomas C. Irving, David A. Kass, Steven Hsu

**Author notes:** **Address Correspondence:** David A. Kass, M.D., Division of Cardiology, Ross Research Building, Room 858, Johns Hopkins Medical Institutions, 720 Rutland Avenue, Baltimore, MD 21205, @dkassjhu, Steven Hsu, M.D., Division of Cardiology, Zayed 7125S, Johns Hopkins Medical Institutions, 1800 Orleans St, Baltimore, MD 21287, @stevenhsu_md.

## Abstract

**Rationale:** Right ventricular (RV) contractile dysfunction commonly occurs and worsens outcomes in heart failure patients with reduced ejection fraction and pulmonary hypertension (HFrEF-PH). However, such dysfunction often goes undetected by standard clinical RV indices, raising concerns that they may not reflect aspects of underlying myocyte dysfunction.

**Objective:** To determine components of myocyte contractile depression in HFrEF-PH, identify those reflected by clinical RV indices, and elucidate their underlying biophysical mechanisms.

**Methods and Results:** Resting, calcium- and load-dependent mechanics were measured in permeabilized RV cardiomyocytes isolated from explanted hearts from 23 HFrEF-PH patients undergoing cardiac transplantation and 9 organ-donor controls. Unsupervised machine learning using myocyte mechanical data with the highest variance yielded two HFrEF-PH subgroups that in turn mapped to patients with depressed (RVd) or compensated (RVc) clinical RV function. This correspondence was driven by reduced calcium-activated isometric tension in RVd, while surprisingly, many other major myocyte contractile measures including peak power, maximum unloaded shortening velocity, and myocyte active stiffness were similarly depressed in both groups. Similar results were obtained when subgroups were first defined by clinical indices, and then myocyte mechanical properties in each group compared. To test the role of thick-filament defects, myofibrillar structure was assessed by X-ray diffraction of muscle fibers. This revealed more myosin heads associated with the thick filament backbone in RVd but not RVc, as compared to controls. This corresponded to reduced myosin ATP turnover in RVd myocytes, indicating less myosin in a cross-bridge ready disordered-relaxed (DRX) state. Altering DRX proportion (%DRX) affected peak calcium-activated tension in the patient groups differently, depending on their basal %DRX, highlighting potential roles for precision-guided therapeutics. Lastly, increasing myocyte preload (sarcomere length) increased %DRX 1.5-fold in controls but only 1.2-fold in both HFrEF-PH groups, revealing a novel mechanism for reduced myocyte active stiffness and by extension Frank-Starling reserve in human HF.

**Conclusions:** While there are multiple RV myocyte contractile deficits In HFrEF-PH, clinical indices primarily detect reduced isometric calcium-stimulated force related to deficits in basal and recruitable %DRX myosin. Our results support use of therapies to increase %DRX and enhance length-dependent recruitment of DRX myosin heads in such patients.

## INTRODUCTION

A third of patients with pulmonary hypertension due to heart failure with reduced left ventricular ejection fraction (HFrEF-PH) develop right ventricular (RV) failure. RV failure is a highly morbid complication of HFrEF-PH, worsening systemic end-organ function, functional capacity, and survival.^1^ However, our ability to identify RV dysfunction remains inadequate as many of these patients go undetected until RV failure is too advanced^2–4^. This raises the possibility that key underlying myocyte contractile defects are not reflected by standard clinical metrics. Prior studies of RV cardiomyocytes extracted from end-stage explanted HFrEF-PH myocardium reported depressed maximum calcium-activated tension^2, 5, 6^ and peak power^5^ although their detection by clinical measures was not reported, nor were biophysical mechanisms that might underlie them. Potential mechanisms include actin-myosin crossbridge kinetics,^7^ sarcomere protein stoichiometry,^8^ thick filament structure and its response to diastolic stretch and load (i.e., length-dependent activation), and the number of myosin heads available for force production. The latter is thought to vary with the proportion of myosin heads residing in a cross-bridge ready state called disordered-relaxed (DRX) versus a super-relaxed (SRX) state which cannot form crossbridges.^9, 10^

Identifying which myofilament-dependent mechanisms contribute to depressed RV contractility in HFrEF-PH has taken on greater importance as pharmaceuticals that stimulate their function are currently being developed and tested for human heart failure. These drugs, known as myotropes, modulate sarcomere proteins to augment contractility without incurring deleterious calcium-mediated metabolic and arrhythmic costs as found with protein kinase A-stimulating inotropes^11^. They each have distinct biophysical effects so optimally leveraging their impact without knowing the underlying myocyte defects is challenging. This may underlie the variable impact of omecamtiv mecarbil, the first myotrope to advance to a large-scale pivotal HF trial, that displayed muted overall benefits^12^.

The present study tested the hypothesis that important features of underlying RV myocyte contractility are not actually reflected by standard clinical parameters of RV function. Determining which are and are not central for developing better more targeted therapeutics. Secondly, we determined biophysical causes for RV contractile defects in HFrEF-PH, focusing on the role of myofibrillar structure, thick-filament activation, and control of the DRX/SRX ratio. The studies were conducted in permeabilized myocytes and muscle strips isolated from RV myocardium from HFrEF-PH patients versus donor controls. Surprisingly, we find that among multiple features reflecting depressed myocyte contractility, only calcium-stimulated isometric tension is reflected in more depressed RV chamber dysfunction defined by clinical indices. Lower peak-activated tension correlates with a lower percent of DRX myosin, reducing association of myosin heads with actin, and with less recruitment of DRX myosin at higher sarcomere length. These data provide new insights into RV myocardial defects in HFrEF-PH, their relation to clinical indices, and the role played by DRX/SRX myosin ratio in this process.

## METHODS

### Study Subjects

RV myocardium was obtained from cardiectomized HFrEF-PH patients (n=23) at time of heart transplantation at the Johns Hopkins Hospital or the Hospital of the University of Pennsylvania. Patients had advanced left ventricular failure (LV ejection fraction, EF < 40%) associated with group II pulmonary hypertension (mean pulmonary artery pressure >20 mm Hg and pulmonary capillary wedge pressure ≥15 mmHg)^13^. Individuals with RV-predominant heart failure (e.g., arrhythmogenic right ventricular cardiomyopathy) were excluded. Non-failing control hearts (n=9) were procured from brain death organ donors whose hearts were not used for transplant primarily due to age. Excised hearts were procured as described^14^ using high-potassium cold-cardioplegia solution and open-chest surgical excision, then placed in cold Krebs-Henseleit Buffer (KHB) on wet ice, and transferred to an on-campus location for rapid dissection, freezing in liquid nitrogen, and storage at −80 °C.

Demographic and basic echocardiographic data obtained just prior to cardiectomy were used. For HFrEF-PH patients, inotropic medications, right heart catheterization hemodynamics and, if available, RV-focused echocardiography were also used for analysis. Investigators acquiring or analyzing clinical data were blinded to myocyte characterizations and vice versa.

### Myocyte Structural and Biomechanical Assays

#### Isolated myocyte studies

Full details are provided in Supplemental Methods. Briefly, permeabilized (skinned) cardiomyocytes were prepared by cutting frozen tissue over dry ice in 5-10 mg pieces and immediately incubating in ice-cold (0°C) isolation buffer with 0.3% Triton X-100 in the presence of protease (Sigma-Aldrich, MO) and phosphatase inhibitors (PhosSTOP, Roche, Germany), as previously described^15, 16^. After washing in isolation buffer with Triton-X100, cardiomyocytes were affixed to a force transducer-length controller (Aurora Scientific, Canada) using ultraviolet-activated adhesive (Norland, NJ). Cells were then transferred into room temperature relaxing buffer. Sarcomere length (SL) was measured by Fourier transformation of digital images (IPX-VGA210, Imperx, FL), adjusted by micro-manipulators (Siskiyou, CA), and set to 2.1 μm.

Isometric tension-calcium relations at a fixed sarcomere length of 2.1 µM, active tension-sarcomere length relations at a fixed calcium of 3.8 µM, resting tension-sarcomere length dependence at a calcium of 0 µM, rate constant of tension redevelopment after acute crossbridge disruption (k_tr_), hyperbolic tension-velocity dependence and from these calculated tension-power relations were determined in the permeabilized myocytes. Details of these measurements and analysis are provided in Supplemental Methods.

In a subset of studies, cardiomyocytes were exposed to 2 µM mavacamten (Selleck Chemicals, Houston, TX) and relaxing and activating solutions made with 5.95 mM deoxy-ATP (a.k.a. dATP, Millipore Sigma, Burlington, MA) instead of ATP; measures were repeated pre- and post-15-minute drug or small molecule exposure.

#### Small Angle X-Ray Diffraction

Full details are provided in Supplemental Methods. Briefly, small angle X-ray diffraction patterns^9^ were acquired at the BioCAT beamline 18ID at the Advanced Photon Source, Argonne National Laboratory, Lemont, Illinois^17^. Flash-frozen tissue pieces (10-15 mg) were cut in pCa 8 relaxing solution and skinned for 1 hour in relaxing solution containing 1% Triton X-100 and 15 mM BDM at room temperature. Muscle was suspended between two hooks in a customized chamber with two Kapton windows in the X-ray path. The preparation was stretched to SL 2.1 μm by monitoring light diffraction patterns from a helium-neon laser (633 nm). X-ray patterns were collected at 2.1 μm SL at pCa 8. In a subgroup, muscle strips were exposed to 2 µM mavacamten (Selleck Chemicals, Houston, TX) and 5.95 mM dATP-containing solutions (Millipore Sigma, Burlington, MA) in lieu of ATP for 15 minutes; and diffraction patterns collected pre- and post-drug exposure. Between 2-4 patterns were collected for each condition per patient, from which equatorial intensity ratio and lattice spacing was acquired. Intensities and spacings of the 1,0 and 1,1 equatorial reflections were measured using the Equator module of the open source MuscleX data analysis package^18^.

#### Myosin ATP Turnover Kinetics

Full details are provided in the Supplemental Methods. The percentage of DRX myosin (%DRX) was determined with a single nucleotide turnover assay^19–21^ applied to skinned single cardiomyocytes. Myocytes were set to 2.1 μm SL in relaxing buffer, washed in rigor buffer (relaxing buffer without ATP or CrP) for 1 minute, incubated for 1 minute in rigor buffer with the fluorescent ATP analog 25 μM 2’-/3’-O-(N’-Methylanthraniloyl) adenosine-5’-O-triphosphate (a.k.a. mant-ATP, Enzo Life Sciences, Axxora LLC, Framingham, NY), and moved to room temperature relaxing buffer. The acquired fluorescence decay was fit to a biexponential to obtain the percentage of DRX myosin. In total, data from >1000 myocytes were acquired.

### Transcriptomic Analysis

MRNA Isolation, sequencing, and informatics are described in Supplemental Methods.

### Unsupervised Machine Learning

Myocyte mechanical properties were used to identify myocyte functional phenotypes. Mechanical features were filtered by their coefficient of variation (ratio of standard deviation to the mean). Features with >50% coefficient of variation were then used for model training. An alternate model including peak power, the slope of the length-tension relationship, and area under the power-tension curve, was also tested. Features for both models were input into an infinite mixture Dirichlet process model.^22^ Hyperparameters were: maximum components, 3; expectation-maximization (EM) tolerance, 1×10^-3^; maximum iterations, 100; weight concentration prior, 10. Models were trained in Python using the sklearn package. Results were visualized with principal component analysis. A third model using clinical characteristics of RV function was also used. Right atrial pressure, the ratio of right atrial-to-pulmonary capillary wedge pressure, pulmonary artery pulsatility index (pulmonary artery pulse pressure/right atrial pressure), tricuspid annular planar systolic excursion, and pulmonary vascular resistance were input into a K Means clustering model (sklearn, Python).

### Spatially Explicit Half-Sarcomere Modeling

Numerical simulations of tension-calcium, tension-power/velocity, and tension-length relationships were performed via FiberSim,^23^ a spatially explicit model of myofilament-level contraction. Thick filament kinetics were simulated as described by Kosta et al.^23^ using a scheme with an SRX state, a DRX state, and a single actin-attached state. The probability of heads transitioning from the SRX to the DRX state increased linearly with force. In addition to control simulations, two other conditions were tested; one in which 25% of myosin heads were confined to the SRX state (unable to recruited to the DRX state), and a second with the same constraint but also with greater proportion (50%) of heads in the SRX state.

### Statistical Analysis

Results are expressed as mean ± SD. Between group comparisons of clinical data were compared with a Mann-Whitney test or Fisher’s exact test for continuous or categorical variables, respectively. Tension-calcium, resting tension, and tension-velocity/power curves were compared using 2-way repeated measure analysis of variance (RM-ANOVA) with Sidak’s multiple comparison test. Fit parameters from these relationships, equatorial intensity ratio, and ATP turnover proportions/rates were compared with a Kruskal-Wallis test with Dunn’s multiple comparisons. For pre- and post-drug studies, tension-calcium relationships were compared with 2-way RM-ANOVA. Delta fit parameters with incubation were compared to zero with a one-sample Wilcoxon test. Analysis was performed using Stata 15.1 or Prism Version 9.0. All MATLAB code used for analysis is publicly available on GitHub (vpjani/myocyte_analysis_code).

## RESULTS

### Patient Characteristics

Demographic, clinical, and histologic characterization of control and HFrEF-PH patient data are provided in **Table 1**. Control and HFrEF-PH patients were matched in age, body mass index, and sex. The etiology of HFrEF-PH was 16% ischemic and 84% non-ischemic, and HFrEF-PH left ventricles were more hypertrophied and fibrotic while RVs were more hypertrophied but had similarly low fibrosis levels as controls. All HFrEF-PH patients were treated with positive inotropes. Additional clinical features of the HFrEF-PH patients are provided in **Table S1**. Primary assessments of RV dysfunction included elevated right atrial pressure (RAP) and volume index, RV end-diastolic dimension, elevated ratio of RAP-to-pulmonary capillary wedge pressure, reduced pulmonary artery pulsatility index, tricuspid annular planar systolic excursion (TAPSE), and global RV and septal strain. Many of these parameters demonstrated substantial inter-subject variability, with coefficients of variance ranging 35%-149% (median 61%). LV dysfunction measures were more consistent.

**Table 1.** Demographics + Clinical Characteristics Control vs. HFrEF-PH.

| Characteristic | Control (n=9) | HFrEF-PH (n=23) | P Value |
| --- | --- | --- | --- |
| Age | 55±11 | 50±12 | 0.225 |
| BMI | 26±5 | 27±6 | 0.628 |
| Gender, % F | 33% (3/9) | 15/25 (60%) | 0.712 |
| DM, % | 0% (0/0) | 33% (6/18) | 0.156 |
| LVMI | 114±24 | 149±43 | **0.026** |
| Creatinine | 0.8±0.3 | 1.2±0.4 | **0.043** |
| **Echocardiographic Characteristics** |  |  |  |
| LV Ejection Fraction | 63±4 | 23±13 | **<0.0001** |
| LV End Diastolic Dimension | 4.1±0.4 | 6.4±1.4 | **0.0008** |
| Posterior Wall Thickness | 0.9±0.1 | 0.9±0.2 | 0.917 |
| **Histologic Characterization** |  |  |  |
| LV, % Fibrosis | 8±2 | 22±10 | **0.001** |
| LV Myocyte CSA, pixels | 1272±390 | 2970±1530 | **0.007** |
| RV, % Fibrosis | 12±7 | 12±5 | 0.413 |
| RV Myocyte CSA, pixels | 1045±248 | 2175±804 | **0.0005** |

BMI – Body Mass Index, DM – Diabetes Mellitus, LV – Left Ventricle, RV – Right Ventricle, LVMI – Left Ventricular Mass Index, CSA – cross sectional area.

### Identification of HFrEF-PH subgroups based on cardiomyocyte mechanics

The pipeline for generation of HFrEF-PH subgroups based on myocyte mechanical measurements is shown in **Figure 1A**. The limited size of our cohort necessitated reducing the number of features used for model training. As such, we constructed the algorithm to include mechanical features with at least 50% greater coefficient of variation in HFrEF-PH versus control myocytes, assuming these were features encoding the most information needed to distinguish subgroups. Four RV myocyte features were agnostically identified for training: T_max_, V_max_, k_tr_ at 3.8 μM [Ca^2+^], and resting tension at 2.6 μm, and used to train an Infinite Process Dirichlet Process model (Stick Breaking Process)^22^. Two clusters of HFrEF-PH patients were identified from this process, with one (blue) being closer to control (black) mechanical properties than the other (red) based on principal component analysis (**Figure 1B)**. The addition of two other key features of cardiomyocyte contractility: cardiomyocyte stiffness at 3.8 μM Ca^2+^ and the area under the tension-power relationship, did not significantly change the cluster composition (Cohen’s Kappa = 0.76), indicating that the model is robust even with modifications in user input.

**Figure 1.**
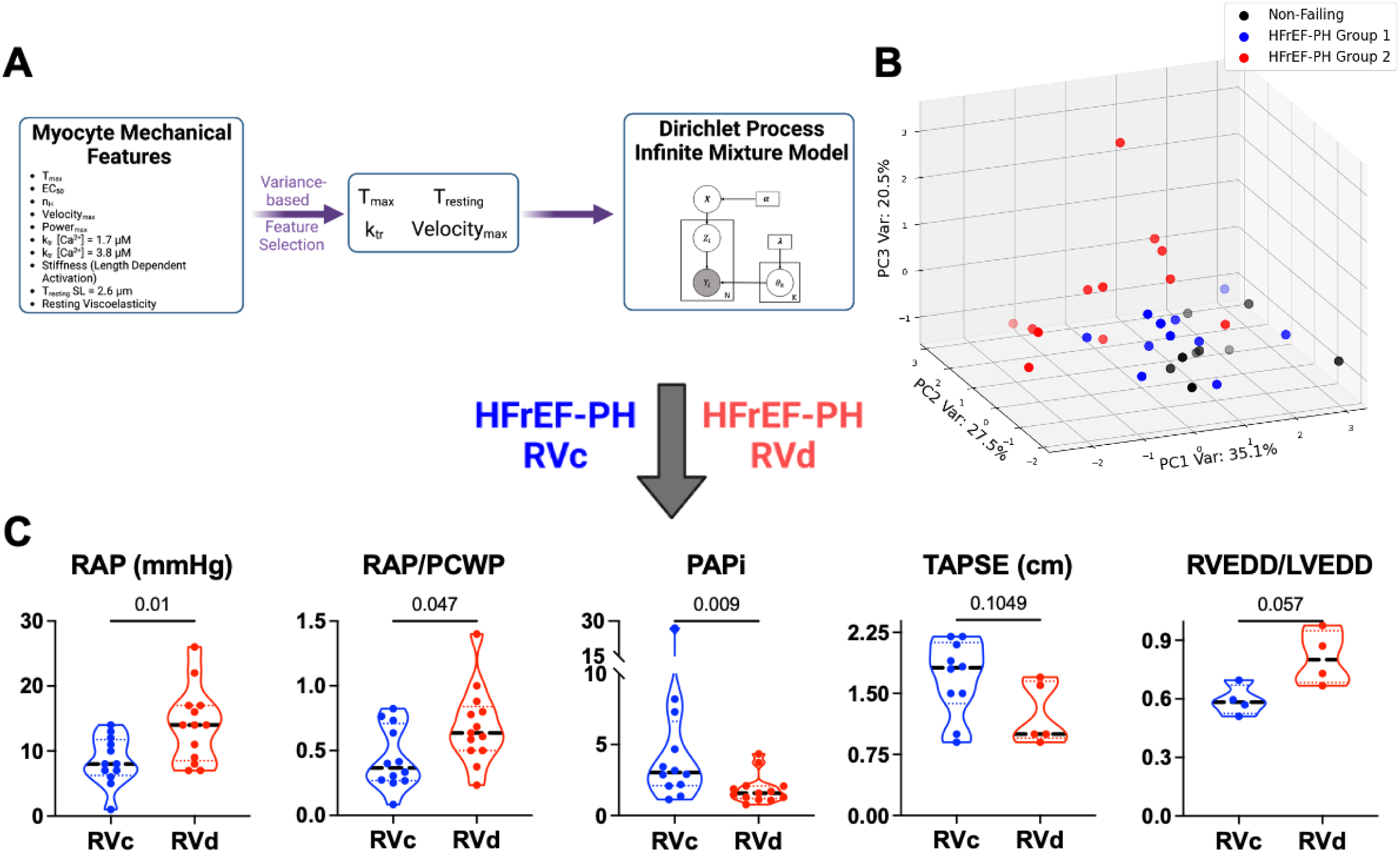
Unsupervised machine learning identifies two sub-groups of cardiomyocyte function in HFrEF-PH. **A)** Schematic of the unsupervised machine learning pipeline. Cardiomyocyte features with coefficient of variation 50% higher than the corresponding coefficient of variation from non-failing cells were incorporated in an infinite mixture Dirichlet process model, and clusters visualized by principal component analysis (PCA). Shading corresponds to depth, with lighter points closer to the origin. **B)** PCA of cardiomyocyte biophysical features (T_max_, V_max_, k_tr_ at 3.8 μM [Calcium], and resting tension at 2.6 μm) included in the model. Three principal components along with corresponding variances are shown. Two HFrEF-PH subgroups were identified: RVc (blue) and RVd (red). The same PCA applied to NF controls is displayed in black. **C)** Comparison of clinical RV hemodynamic features in RVc versus RVd groups. RAP – right atrial pressure, RAP/PCWP – RAP to pulmonary capillary wedge pressure ratio, PAPi – pulmonary artery pulsatility index, TAPSE – tricuspid annular planar systolic excursion, and RVEDD/LVEDD – Right-to-left ventricular end diastolic dimension ratio. P-values are from Mann-Whitney test. RVc – Right Ventricular compensation. RVd – Right Ventricular decompensation.

We next compared the clinical features of RV function between the two myocyte-mechanics derived subgroups (**Figure 1C**) and found one group had worse RV function (RVd) than the other (RVc), including lower pulmonary artery pulsatility index (PAPi), higher right atrial pressure and ratio to pulmonary capillary wedge pressure, borderline increased RV to LV dimension ratio. TAPSE, a common RV functional metric was not significantly different.

### Isometric Ca^2+^-activated tension discriminates HFrEF-PH subgroups

**Figure 2A** displays isometric tension-calcium relations for control, RVc, and RVd myocytes. All groups had similar tension at low (diastolic range) Ca^2+^, but at higher (systolic) levels (e.g. ≥ 2.5 mM Ca^2+^), isometric tension was markedly depressed in RVd yet remained near control in the RVc group. EC_50_ was similarly reduced in both groups, indicating increased calcium sensitivity independent of the clinical metrics (**Figure S1A**). Cooperativity (Hill coefficient, **Figure S1B**), resting length-tension relationships (**Figure S1F)**, and passive titin-based viscoelasticity were also all similar between groups (**Figure S1G**).

**Figure 2.**
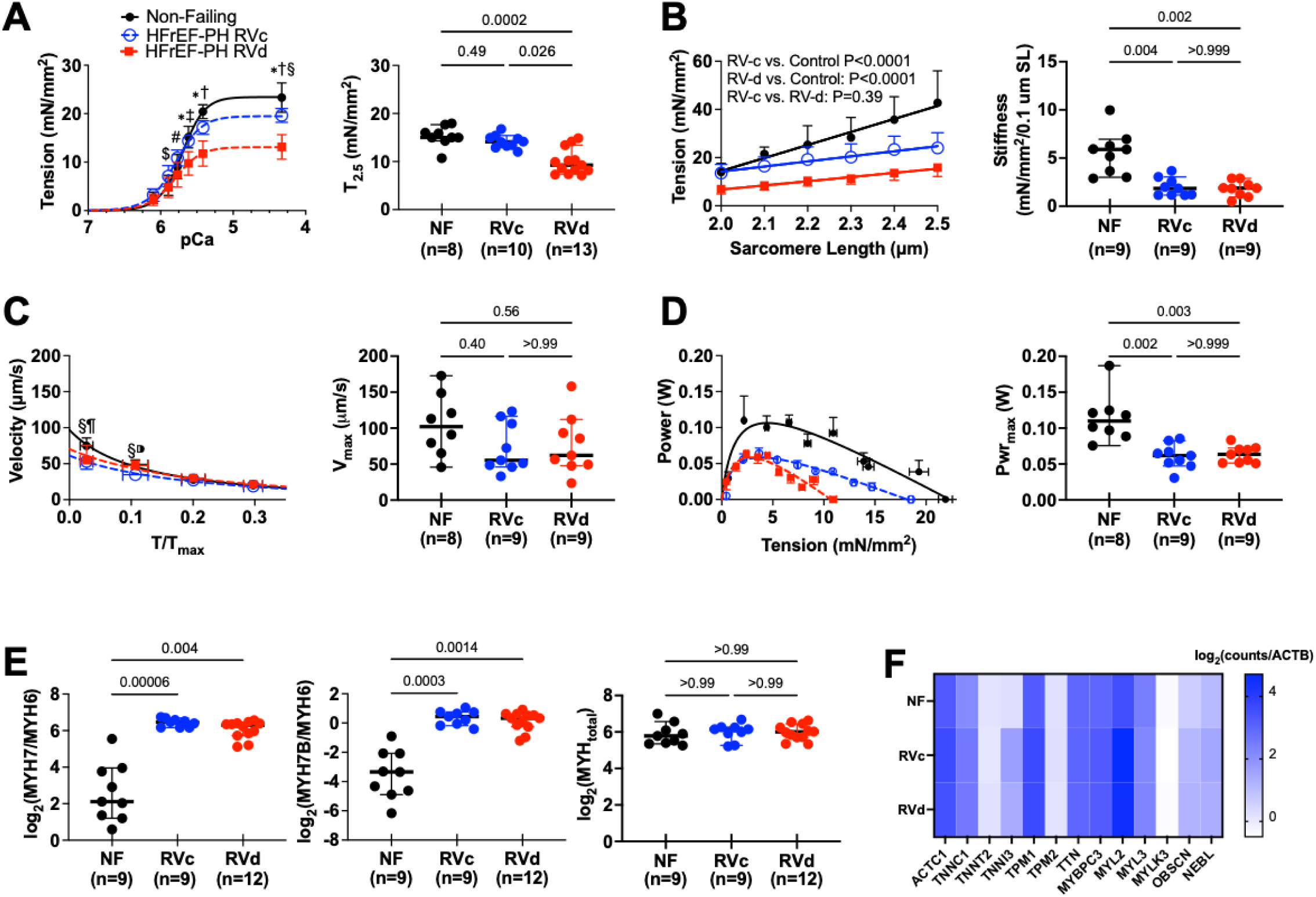
Isometric tension, but not other measures of cardiomyocyte contractility, associate with clinical indices of RV function. **A)** Tension-calcium relationships (mean±SD) for each HFpEF-PH subgroup as defined by machine learning compared to controls (black). 2WANOVA: P=3.9e-7 for group effect, and <1e-13 for Ca^2+^ dependence, and Ca^2+^-group interaction. Symbols for post-hoc Sidak multiple comparison test (P= $ 0.01, # 0.006, * ≤7e-8 RVc vs RVd groups; † <1e-13, ‡ 4e-9 RVd vs non-failing (NF); and § 9e-4 RVd vs Control). **B)** Tension-sarcomere length relationships measured at activating Ca^2+^ for each patient. P-values are results of group pair-wise analysis of covariance for a difference in slope (e.g. active myocyte stiffness). **C)** Absolute velocity-normalized tension relationship for each subgroup. P-value reflects group effect by 2-way-RMANOVA, *Sidak multiple comparison test at each normalized load. RVd vs. NF: ¶ P=0.004; RVc vs. NF: §P<0.0037; RVd vs. RVc: ⁍P=0.006. **D)** Absolute power-absolute tension relationship for each subgroup. RVc-RV compensated. RVd – RV decompensated. **E) Left to right:** Ratio of β-myosin (MYH7)/α-myosin (MYH6), MYH7B/α-myosin, and total myosin heavy chain (MYH_total_) isoform gene counts for three groups. P-values from Dunn’s Multiple comparisons after a Kruskal-Wallis test. **F)** Heat map for sarcomere protein genes generally shows very consistent expression levels in all three patient groups.

Surprisingly, other primary measures of myocyte contractility were similarly reduced in both RVc and RVd subgroups. One was the slope of the calcium-activated tension-length relationship, a myocyte-level correlate of chamber end-systolic stiffness (e.g., elastance) (**Figure 2B**). Both groups had equally depressed slopes, though unlike RVc, the RVd relation was further shifted downward, consistent with the lower T_2.5_ in this group. Furthermore, maximal unloaded shortening velocity (V_max_), maximal power (Pwr_max_), and tension at peak power, all primary indices of contractile function, were similarly reduced in both groups (**Figure 2C, 2D, Figure S1C**). Kinetic measures of crossbridge formation, including rate of tension redevelopment (k_tr_) were not significantly different between any groups (**Figure S1D, S1E, Table 2)**. Importantly, myocyte cross-sectional area and cell length were similar between groups (**Figure S2**), so altered cell geometry was not a significant factor in these results.

**Table 2.** Myocyte Mechanical Parameters in the two HFrEF-PH subgroups (RVc, RVd) identified by Infinite Mixture Dirichlet Process Modeling. P values are from Mann-Whitney test. T_2.5_– Isometric calcium activated tension at 2.5 μM calcium, nH – Hill Coefficient, EC_50_ – calcium concentration required for half-maximal activation, V_max_ – maximum unloaded shortening velocity, Power_max_ – maximum power generation, k_tr_ – constant for crossbridge reformation, T_resting_ – Resting tension, *τ*_*σ*_ – time constant for creep in strain, *τ*_*ϵ*_ – time constant for stress recoil. T_opt_ – Tension at maximum power.

|  | HFrEF-PH<br>RVc (n=10) | HFrEF-PH<br>RVd (n=13) | P-Value<br>(RVc vs. RVd) |
| --- | --- | --- | --- |
| **Mechanical Properties** |  |  |  |
| $T_{2.5}$ (mN/mm<sup>2</sup>) | 14 ± 1 | 9 ± 3 | **0.003** |
| nH | 2.5 ± 0.4 | 2.6 ± 0.6 | 0.66 |
| $EC_{50}$ | 1.5 ± 0.2 | 1.6 ± 0.2 | 0.46 |
| $V_{max}$ (m/s) | 53 ± 15 | 55 ± 36 | 0.97 |
| $Power_{max}$ (W) | 0.06 ± 0.02 | 0.06 ± 0.01 | 0.76 |
| $T_{opt}$ (mN/mm<sup>2</sup>) | 4.0 ± 1.5 | 2.8 ± 0.8 | 0.06 |
| Tension-Power (W mN/mm<sup>2</sup>) | 0.56 ± 0.13 | 0.36 ± 0.09 | **0.007** |
| $k_{tr}$ (s<sup>-1</sup>), [Calcium] = 1.7 μM | 0.8 ± 0.4 | 1.0 ± 1.5 | 0.20 |
| $k_{tr}$ (s<sup>-1</sup>), [Calcium] = 3.8 μM | 1.2 ± 0.7 | 1.0 ± 0.5 | 0.83 |
| Stiffness, [Calcium] = 3.8 μM<br>(mN/mm<sup>2</sup>/0.1 m) | 2.0 ± 0.9 | 1.8 ± 0.8 | 0.83 |
| $T_{resting}$ (mN/mm<sup>2</sup>), SL = 2.6 m | 7.5 ± 1.2 | 7.4 ± 2.8 | 0.66 |
| $\tau_{\sigma}$ (s) | 1.4 ± 0.2 | 1.7 ± 1.1 | 0.69 |
| $\tau_{\epsilon}$ (s) | 0.6 ± 0.3 | 0.4 ± 0.2 | 0.30 |
| **Clinical Data** |  |  |  |
| Inotropes (Milrinone/Dobutamine) | 10 (100%) | 13 (100%) | 1.00 |
| Right Atrial Pressure, mmHg | 9 ± 4 | 14 ± 6 | **0.01** |
| Pulmonary Artery Systolic Pressure, mmHg | 48 ± 10 | 45 ± 14 | 0.46 |
| mean Pulmonary Artery Pressure, mmHg | 29 ± 8 | 30 ± 10 | 0.76 |
| Pulmonary Capillary Wedge Pressure, mmHg | 21 ± 7 | 22 ± 8 | 0.44 |
| Pulmonary Artery Pulsatility index | 5.5 ± 7.1 | 1.9 ± 1.0 | **0.009** |
| Right Atrial Pressure/Pulmonary Capillary Wedge Pressure | 0.4 ± 0.2 | 0.7 ± 0.3 | **0.047** |
| Cardiac Output, L/min | 4.0 ± 1.8 | 4.1 ± 1.1 | 0.42 |
| Cardiac Index, L/min/m<sup>2</sup> | 2.0 ± 0.9 | 2.2 ± 0.6 | 0.23 |
| Stroke Volume, mL | 50 ± 27 | 73 ± 57 | 0.12 |
| Pulmonary Vascular Resistance, Wood units | 2.8 ± 1.6 | 2.2 ± 1.2 | 0.23 |
| Tricuspid Annular Planar Systolic Excursion, cm | 1.7 ± 0.5 | 1.2 ± 0.4 | 0.10 |
| RV End Diastolic Dimension, cm | 4.0 ± 0.7 | 4.0 ± 0.4 | 0.72 |
| LV Ejection Fraction, % | $22 \pm 10$ | $25 \pm 18$ | 1.00 |
| LV End Diastolic Dimension, cm | $6.6 \pm 1.4$ | $5.8 \pm 1.5$ | 0.30 |
| Right Atrial Volume, mL | $108 \pm 12$ | $211 \pm 67$ | **0.03** |
| Right Atrial Volume index, mL/m<sup>2</sup> | $54 \pm 13$ | $115 \pm 51$ | **0.04** |
| Fractional Area Change, % | $26 \pm 5$ | $25 \pm 5$ | 0.48 |

To further test the robustness of the mapping between clinical indices and myocyte mechanics, the model process was reversed using hemodynamic and echocardiographic RV function measures to define the subgroups and then contrasted their myocyte properties. This yielded the same result as the model above, with only T_2.5_ different between HFrEF-PH sub-groups (**Figures S3, S4**). This confirms that the relationship between resting hemodynamics and isometric calcium-stimulated tension is monotonic.

### Myofilament protein stoichiometry and myosin isoform in HFrEF-PH

Myocyte ensemble force (F_ens_) is given by the equation:

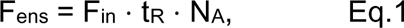

where F_in_ is the intrinsic force of the sarcomere, t_R_ is the duty ratio (time myosin is bound to actin during the crossbridge cycle), and N_A_ is the number of myosin heads available for force production.^24^ We explored each of these determinants to identify potential factors underlying the decline in T_2.5_ in RVd myocytes. An important mediator of F_in_ is the stoichiometry of sarcomere components and number of myofibrils within cardiomyocytes, both being transcriptionally regulated.^8^ We therefore performed bulk RNA-seq on RV tissue. Pathway analysis of HFrEF-PH versus controls revealed upregulated striated muscle differentiation and sarcomere cytoskeleton, and downregulated metabolism pathways (**Figure S5**), consistent with prior reports in HFrEF patients^25^. While 933 genes were differentially expressed between control and HFrEF-PH (**Figure S6A**), there were no significant differences detected between HFrEF-PH subgroups (**Figure S6B**). Thus, altered sarcomere component stoichiometry was unlikely a factor for HFrEF-PH subgroup F_ens_ differences.

The duty ratio t_R_ in Eq. 1 reflects the relative time per unit interval with active crossbridge engagement and so impacts crossbridge kinetics. One factor that can influence such kinetics is the myosin isoform^26, 27^. Both beta/alpha myosin heavy chain ratio (MYH7/MYH6, **Figure 2E**) and MYH7B/alpha-myosin heavy chain ratio were similarly elevated in the HFpEF-PH subgroups. As both MyH7 and MYH7B have slower kinetics^28^, this could contribute to depressed Pwr_max_ and V_max._ in both groups. Total myosin expression was the same in all three groups (**Figure 2E**), and no differences in gene expression of other sarcomere proteins were found (**Figure 2F**).

### HFrEF-PH subgroups display different myofilament structure

To determine if RVd myocytes had fewer myosin heads functionally available for force generation (e.g., reduced N_A_ in Eq. 1), small angle X-ray diffraction patterns were obtained from isolated skinned RV muscle strips. When X-rays pass through the semi-crystalline sarcomere structure, they yield diffraction patterns (**Figure 3A**), with the 1,0 equatorial reflection intensity, I_1,0_, defined by thick filament densities and the 1,1 equatorial reflection intensity, I_1,1_, by a combination of thick and thin filament densities (**Figure 3B**). The ratio of their intensities, the equatorial intensity ratio (I_1,1_/I_1,0_), correlates with the proximity of myosin heads to the thin filaments under relaxing conditions ^29^. During contraction, I_1,1_/I_1,0_ correlates with the number of actin-myosin cross bridges formed and subsequent force development.^29, 30^ Under relaxing conditions, reduced I_1,1_/I_1,0_ indicates decreased association between myosin heads and thin filaments and more myosin heads associated with the thick filament backbone. This reflects thick filament inactivation and reduced availability of myosin for force generation.

**Figure 3.**
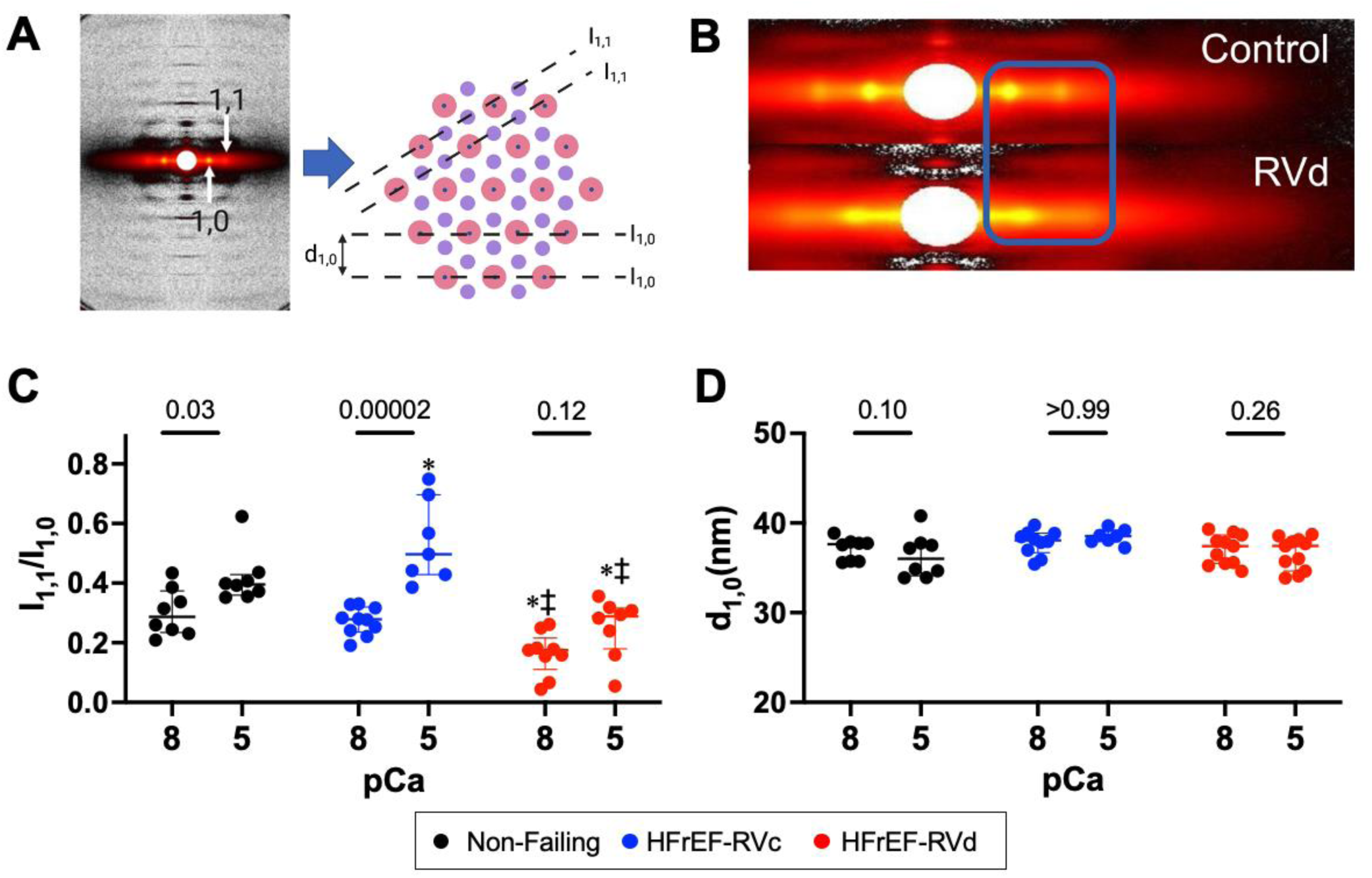
The thick filament is inactivated in HFrEF-PH RVd patients. **A)** Example X-ray diffraction pattern from frozen human myocardium and schematic of how (1,1) reflecting thick-thin filament engagement, and (1,0) reflecting thick filaments based on their equatorial intensities are quantified. Intensities were obtained by averaging a 2D window in MuscleX v1.5 into a 1D intensity profile. **B)** Representative X-ray diffraction patterns showing the equatorial intensity ratio shift with decreased magnitude of I_1,1_ in HFrEF-PH RVd patients vs. controls. **C) *Left:*** Equatorial intensity ratio, I_1,1/_I_1,0_, for NF (black) and HFrEF-PH RVd (blue) and RVd (red) at: pCa = 8 (relaxing conditions) and pCa = 5 (maximum activation). ***Right:*** Lattice spacing (d_1,0_) quantified from the equators of X-ray diffraction patterns for control (black), HFrEF-PH RVc (blue), and HFrEF-PH RVd (red) at pCa 8 (relaxing) and pCa 5 (maximum activation). All diffraction patterns acquired at 2.1 μm sarcomere length. P-values are from 2-way repeated measures ANOVA. *P<0.03 vs. non-failing. ‡P<0.007 vs. HFrEF-PH RVc.

X-ray data revealed that I_1,1_/I_1,0_ is depressed in HFrEF-PH RVd but not RVc, both compared to non-failing controls (**Figure 3C**). Depressed I_1,1_/I_1,0_ in RVd under relaxing conditions remained so when the fibers were placed in maximal calcium activating solution (10 µM, pCa=5, **Figure 3C**). Beyond calcium binding to troponin to activate contraction, calcium-dependent structural activation of the thick-filament has also been reported^31^. We found increased I_1,1_/I_1,0_ ratio in control and RVc groups when Ca^2+^ was increased, but this did not occur in RVd (2W-ANOVA, Ca^2+^-Group Interaction p=0.01). Another factor that can depress I_1,1_/I_1,0_ is a decline in the spacing between the thick filaments or lattice spacing. However, this was not significantly different in relaxing or activating conditions among the three groups (**Figure 3D**). Together, these data support a reduction in the number of available myosin heads (lower N_A_) in HFrEF-PH RVd patients as a primary mechanism for reduced ensemble averaged myocyte force (tension). They also show that calcium-dependent activation of the thick filament^31^ is impaired in patients with depressed calcium-stimulated isometric tension.

### The proportion of DRX myosin is lowest in the HFrEF-PH group with depressed T_2.5_

Differences in I_1,1_/I_1,0_ between the two HFrEF-PH groups could translate to a disparity in the proportion of myosin heads residing in the DRX state.^7, 9^ This was tested by measuring ATP turnover rate with fluorescent mant-ATP in skinned RV cardiomyocytes. The mant-ATP signal decay is fit to a biexponential decay, and SRX myosin identified by secondary slower fluorescence decay (T_2_) following an initial rapid decay (T_1_). T_1_ is thought to relate to acute ATPase, release of non-specifically bound nucleotides, and diffusion of nucleotides from the cell. Thus, proportions of SRX and DRX myosin are quantified from the slow phase and in relaxing conditions are considered to add up to 100%. **Figure 4A** shows that in RVd, a larger proportion of fluorescence decay in the mant-ATP assay occurs during the slow late phase indicating reduced %DRX (31±23% versus 51±15% for control, P=0.001). In contrast, %DRX in RVc is similar to control (43±23%, p=0.1, **Figure 4B**). Both T_1_ (**Figure 4C**) and T_2_ (**Figure 4D**) decay constants were similar in RVd vs control, whereas T_1_ was lower in RVc vs. control. Lastly, we find the %DRX directly correlated with T_2.5_ (P=0.003 overall, P=0.03 after adjusting for patient group as covariate, **Figure 4E**), further supporting their association.

**Figure 4.**
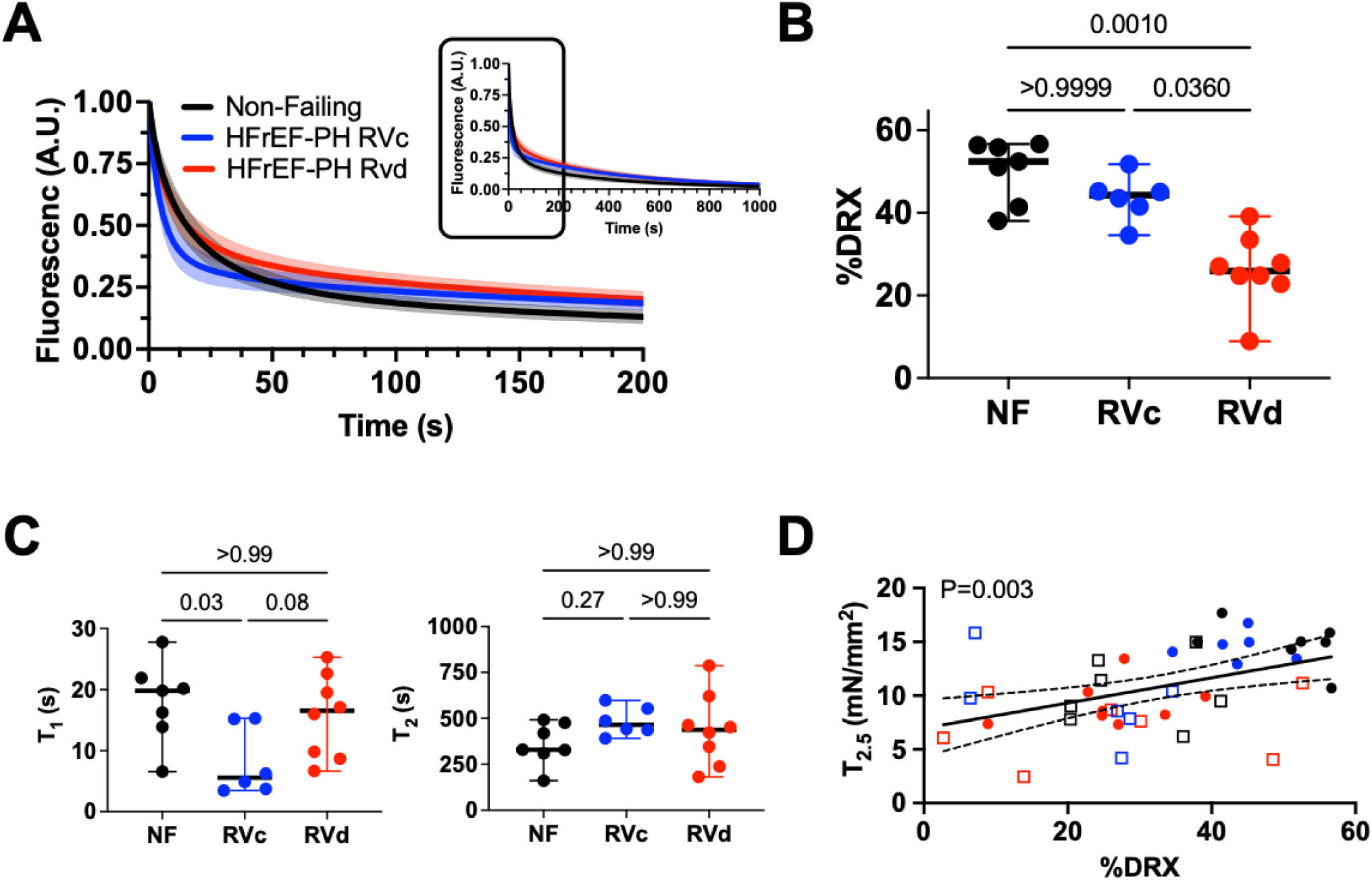
Proportion of disordered-relaxed (DRX) myosin is reduced in HFrEF-PH RVd patients. **A)** Group averaged bi-exponential fits derived from fluorescent turnover assay (mant-ATP). Mean (center bold line) and 95% confidence intervals shown for each group. Upper insert shows full time course. **B)** Summary data for %DRX and **C)** fast (T_1_) and slow (T_2_) ATP turnover rates for each group. P-values for group effects are from general linear models that adjust for individual cardiomyocytes, the patient, and their interaction. Each dot reflects mean result for a given patient. **D)** Linear regression between %DRX and isometric tension (T_2.5_∼ %DRX + Group) at 2.5 µM Ca_2+_ (T_2.5_) for all three groups, including baseline (colored circle) and mavacamten treated (open square) conditions.

### Change in %DRX differentially alters isometric active tension in HFrEF-PH subgroups

To further explore the relationship between %DRX and isometric Ca^2+^-dependent tension, myocytes were exposed to either a myosin inhibitor (mavacamten) that reduces %DRX or 2’-deoxy-ATP (dATP) that increases it.^9, 32–34^ In RVc myocytes, mavacamten reduced Ca^2+^-tension dependence over a broad range (**Figure 5A**), whereas this impact was much less in RVd myocytes. To the right are the I_1,1/_I_1,0_ ratios measured by X-ray diffraction of myocardium from the same patients (± mavacamten) that shows reduction in the ratio in RVc but not RVd myocytes. The opposite responses were obtained with dATP that increases %DRX; there was little change in Ca^2+^-activated tension or I_1,1/_I_1,0_ ratio in RVc myocytes but a significant increase of both in RVd (**Figure 5B**). These data show that myocytes with less DRX myosin at baseline have greater response to agents that increase it (e.g., RVd) than in those where it is close to normal (RVc), whereas agents that increase %DRX are more impactful in the latter group. This highlights a need for more personalized approaches to using myotropes.

**Figure 5.**
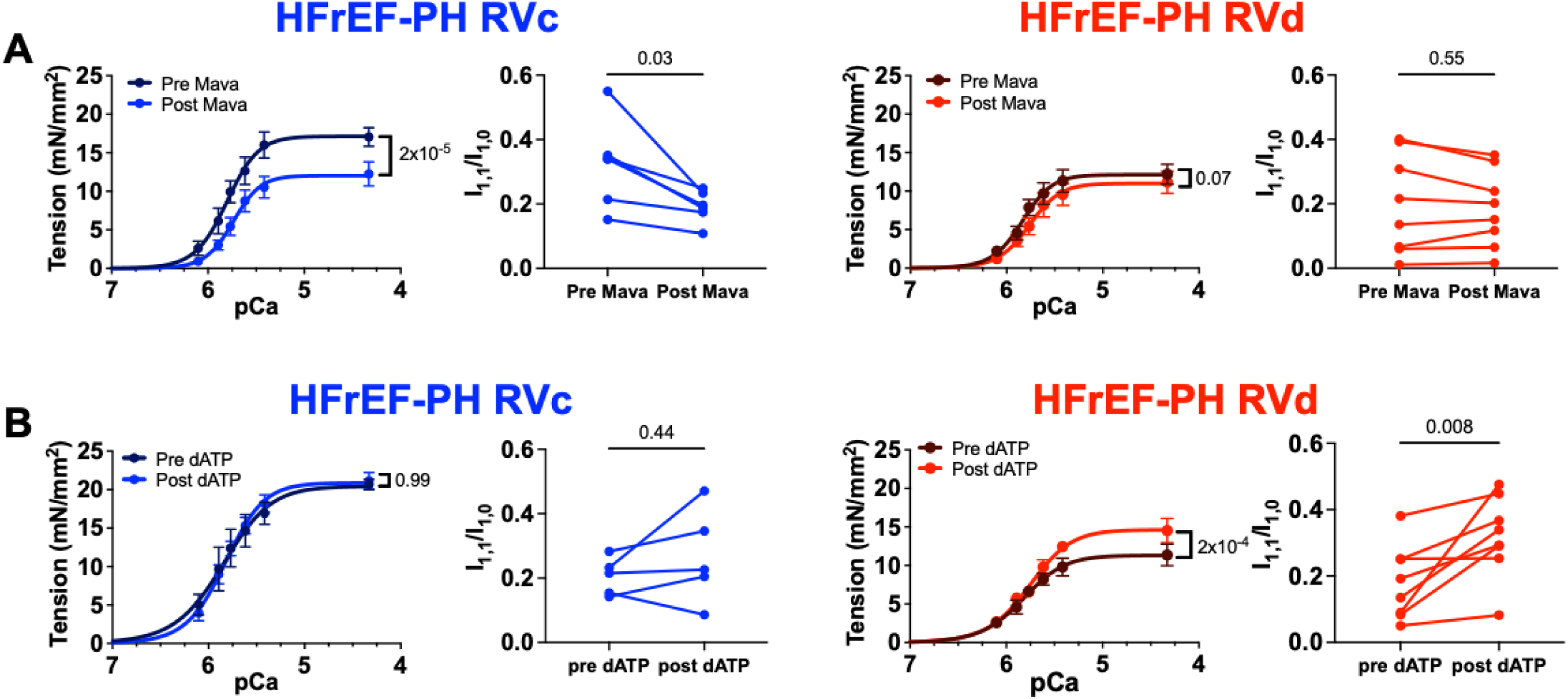
Pharmacologic manipulation of the proportion of DRX myosin in HFrEF-PH RVc and RVd. **A)** Tension-calcium relationship pre- and post-mavacamten (2 μM) for ***left:*** HFrEF-PH RVc and ***right:*** HFrEF-PH RVd. P-value reflects the group effect by 2-way-RMANOVA. *p<0.05 pre vs. post incubation, Sidak multiple comparison test. Individual I_1,1/_I_1,0_ obtained from skinned cardiomyocyte strips from ***left:*** RVc and ***right:*** RVd are shown to the right of their respective tension-calcium relationships. P-values are paired Wilcoxon tests. **B)** Tension-calcium relationship pre- and post-dATP (5 mM) for ***left:*** HFrEF-PH RVc and ***right:*** HFrEF-PH RVd. P-value reflects the group effect by 2-way-RMANOVA. *p<0.05 pre vs. post incubation, Sidak multiple comparison test. Individual I_1,1/_I_1,0_ obtained from skinned cardiomyocyte strips from ***left:*** RVc and ***right:*** RVd are shown to the right of their respective tension-calcium relationships. P-values are paired Wilcoxon tests.

### Myocyte length-dependent tension and strain-dependence of %DRX myosin

To identify mechanisms for depressed myocyte length-dependent tension, we used a spatially explicit half sarcomere model based on the opensource program FiberSim^23^, assuming a three-state kinetic model for thick filament activation. Two other conditions were tested; both prevented recruitment of SRX heads to DRX, with one further increasing the percent of SRX. The last condition best replicated the experimentally observed tension-calcium, tension-length, and tension-power/velocity relationships (**Figure 6A**, c.f. **Figure 2**), indicating that both a greater %SRX and impaired SRX to DRX transition were required. Our prior experiments confirmed the former. To test the later, SRX/DRX balance was determined by mant-ATP assay before and after increasing sarcomere length. We found control cardiomyocytes when stretched from 2.1 to 2.4 μm SL in relaxing conditions displayed a 51% rise in %DRX (**Figure 6B**) but this was substantially depressed (22%) in myocytes from either HFrEF-PH subgroup. These data provide a novel mechanism for depressed myocyte active stiffness showing reduced strain-dependent recruitment of DRX myosin plays a role.

**Figure 6.**
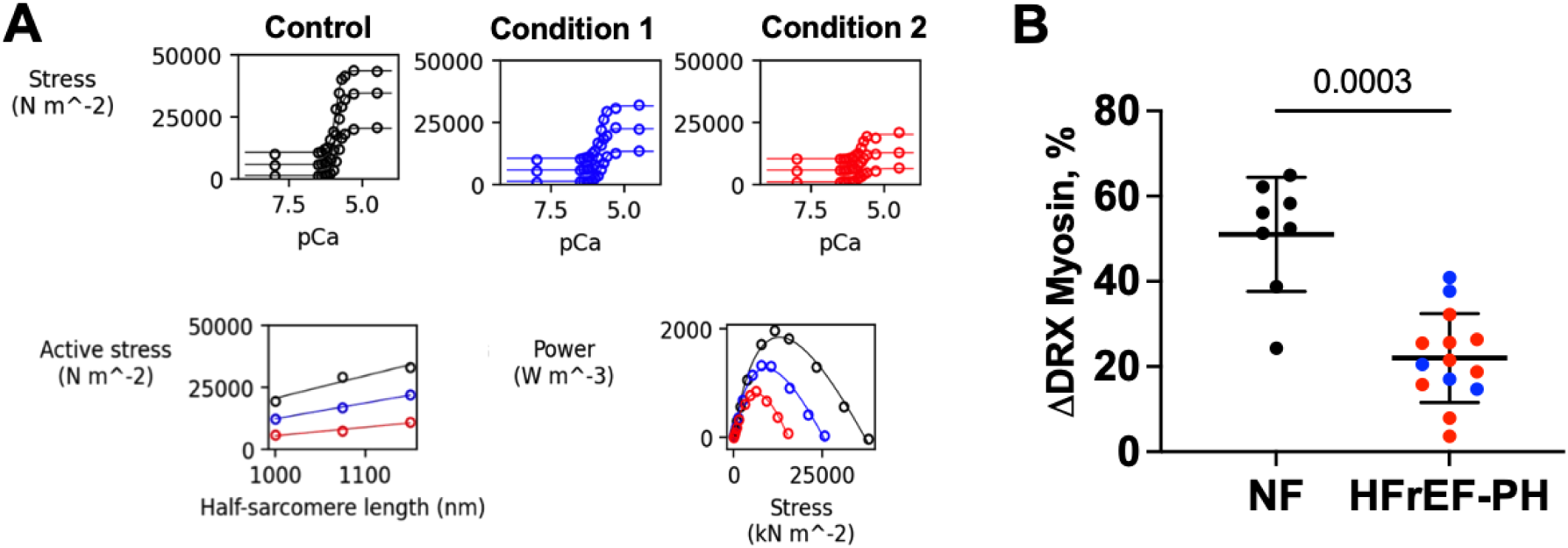
Inactivated myosin in both HFrEF-PH subgroups is not as sensitive to length-mediated recruitment. **A)** Results from FiberSim simulation in three conditions, a control state (black), a condition where 25% of the myosin are confined to the SRX state and cannot transition to the DRX state (Condition 1 - Blue), and one having this same feature but with a higher proportion of SRX heads (50%) in this state (Condition 2 - Red). ***Top:*** Isometric tension-Ca^2+^ relations obtained at three different half-sarcomere lengths (1000 nm [low], 1075 nm [middle], and 1175 nm [high]); ***Lower left:*** Active length-tension relations for each condition (passive tension at each half-sarcomere length was subtracted off); and ***Lower Right***: absolute tension-absolute power relations for each condition. To achieve both reduced peak power and decline in active sarcomere stiffness, a higher proportion of myosin confined the SRX state was required, suggesting a population of SRX myosin heads not able to be recruited to DRX is present in all HFrEF-PH patients and that this is enriched in those patients with RVd. **B)** This prediction was then tested using mant-ATP fluorescence decay curves at sarcomere length of 2.1 versus 2.4 μm. %DRX increased in controls, but significantly less in both HFrEF-PH groups (Mann-Whitney test comparing controls to combined HFrEF-PH; HFrEF-PH RVc: blue, HFrEF-PH RVd: red.

## DISCUSSION

One of the main aims of this study was to determine which fundamental RV myocyte mechanical abnormalities present in HFrEF-PH myocardium are reflected in standard clinical indices of RV function. We surprisingly found only isometric Ca^2+^-stimulated active tension and associated deficits of %DRX myosin associate with clinical RV function indices in these patients. Furthermore, calcium-dependent activation of the thick filament is also only impaired in patients with depressed isometric active tension. Other major mechanical defects, including reduced Pwr_max_, V_max_, myocyte active stiffness, myosin isoform switching, and length-mediated augmentation of %DRX are abnormal in HFrEF-PH patients more generally.

The second goal was to identify biophysical mechanisms for the abnormal RV myocyte contractile behavior found in HFrEF-PH. Our study reveals depressed calcium-stimulated isometric tension that correlates with reduced %DRX myosin and calcium-dependent thick filament activation. To our knowledge, these are the first data revealing such behavior in human hearts with depressed contractility and support a broad relevance of the DRX/SRX balance^36^ to contractility in heart disease. Second, we show that increasing sarcomere length (preload) raises %DRX, likely contributing to the positive slope of the active length-tension relation. Both features are depressed in HFrEF-PH. The decline in length-dependent %DRX has not been shown in HF syndromes previously. These new results provide a roadmap for ongoing efforts to design myotropes for human HF syndromes, revealing specific features to be targeted and central involvement of %DRX.

### Coupling clinical RV assessment to myocyte mechanics

The global disparity between myocyte and clinical manifestations of RV disease in the current study was somewhat surprising. Essentially the overall population of HFrEF-PH RV myocytes have depressed active stiffness (analogous to low chamber end-systolic elastance) and peak power, indicating that something else appears to compensate RV function in HFrEF-PH RVc patients that is not present in RVd patients. One possibility is accelerated ATPase activity of force producing DRX myosin or preserved calcium-dependent activation of the thick filament, and both hypotheses warrant further testing. Our study also leaves open the potential for calcium-cycling adaptations that might counter more coherent depression of sarcomere-force generation. This analysis requires fresh tissue and cells and would not have been practical for the type of measurements (e.g., X-ray diffraction studies) done here. Part of the mystery may be that our clinical indices are simply too coarse or integrated to reflect this myocyte dysfunction. Some evidence supporting this possibility is found in a prior study of RV disease in patients with primary PH with or without systemic sclerosis. In this instance, clinical indices were generally similar between the groups, yet pressure-volume analyses revealed depressed end-systolic elastance and other contractility markers only in those with systemic sclerosis^15^. Use of similar measures in the HFrEF-PH RV has yet to be performed, but it might improve our ability to identify abnormalities at the myocyte level.

### SRX/DRX balance and role in depressed RV myocyte function in HFrEF-PH

Myosin heads are thought to exist in either DRX or SRX states with only DRX myosin effectively interacting with actin to undergo crossbridge cycling in the presence of calcium^36^. Our results support that SRX myosin can be recruited to DRX with a rise in calcium or strain to increase force, and %DRX during activation in turn influences the maximum number of crossbridges and thus tension development with calcium activation^32^. The correlation between active isometric tension and basal %DRX and concurrent depression in HFrEF-PH patients with greatest RV dysfunction is important. It suggests these behaviors have high impact on the disease, as these same clinical indices predict poor clinical outcomes^4^. This also supports investigations of drugs to increase %DRX and enhance active Ca^2+^-dependent myofibrillar tension in RVd patients. The structural changes found in RVd which likely translate to less crossbridges forming during maximum activation also reveal impaired calcium-responsiveness of the thick filament, a relatively new concept of contractility modulation^24, 31, 32, 37^. The greater rise in I_1,1_/I_1,0_ ratio and Ca^2+^-stimulated tension by dATP in RVd versus RVc myocytes is consistent with their differences in basal %DRX.

Our results do not provide a precise biochemical mechanism by which the DRX/SRX balance is altered in HFrEF-PH; however, this was not our goal. Indeed, the precise causes of altered DRX/SRX balance be they induced by calcium or length activation, post-translational modifications, or changes in sarcomere proteins such as mutations are still largely unknown. Even mutations causing hypertrophic cardiomyopathy that cause an increase in %DRX do so by mechanisms that remain to be determined. Prior studies have reported that modification of both myosin binding protein C^19^ and myosin light chain^38^ by phosphorylation is associated with an increase in %DRX. However, blocking their modification as with phospho-site mutations does not lower %DRX, making it less likely these are relevant regulators for HFrEF-PH^19^. An intrinsic limitation of this analysis is that manipulations to alter %DRX do more than that in the myocyte since there is no known specific factors to engage, and this makes cause and effect analyses difficult.

The increase in %DRX at higher sarcomere lengths and its depression in HFrEF-PH provides a novel link between length-dependent activation and availability of myosin to form crossbridges. Since titin stiffness was adjusted for and the influence of calcium sensitivity excluded, the slope of the length-active tension relations measured here relates to two primary mechanisms: strain-dependent activation of the thick filament and increased stiffness of crossbridges. Of these, the first is dependent on %DRX,^39^ and so basal deficits in %DRX may also be expected to directly correlate with the length-tension slope. Yet, this was not observed. However, reduced strain-mediated increase in %DRX was found and correlated with depressed length-tension slope, and this may be the more important feature of DRX myosin relevant to active muscle stiffness. %DRX (and tension) measured at low strains associates with the y-axis intercept of the length-tension relationship. This is consistent with prior studies where decreasing %DRX with mavacamten induced a downward shift in the length-tension relation but no change in slope^9^, and increasing %DRX with dATP also has had no impact on the slope^7^. By contrast, depression of strain-mediated %DRX recruitment would impact the slope. Interestingly, this mechanism has not been observed in rat myocardium^40^ but has been in porcine^9^ and as shown here, in human myocardium. The cause for such differences, perhaps related to sarcomere protein isoforms remains to be determined, but the data suggest use of larger mammalian models may be advisable to better predict human drug responses.

### Study Limitations

Myocardium was obtained from end-stage HFrEF-PH so the generalizability of the results to earlier disease timepoints remains to be determined. Not all clinical or myocyte indices were acquired in all patients primarily for reasons related to tissue availability, which could introduce Type-II errors. Resting length-dependent measures were assessed in the absence of 2,3-butanedione monoxime (BDM), and so neglected the influence of stochastic cross-bridge formation and weak binding on diastolic force. However, this was not relevant to systolic tension measurements as rest tension was adjusted for. The %DRX myosin by mant-ATP assay was only obtained under relaxing conditions and not maximum activation. However, we found with a single cell fluorescent ATP assay, the signal decline with accelerated myosin ATPase activity from Ca^2+^-activation lowered the signal/noise ratio that made analysis impossible. X-ray diffraction patterns mitigate this concern as equatorial intensity ratio was depressed during maximum activation. Likewise, quantification of tension-mediated recruitment of DRX myosin was only acquired in relaxing conditions with diastolic stretch, so effects of calcium were not considered. As noted, post-translational modifications that may underlie a basal increase in DRX myosin were not identified, however, none have yet been confirmed and this will undoubtedly take considerable future efforts to achieve.

## CONCLUSIONS

The dissociation of underlying myocyte contractile defects with standard clinical RV function metrics supports the need for more other specific indices of RV function in HFrEF-PH, and likely all HF patients. That clinical indices do identify the subgroup of patients with depressed isometric calcium-activated tension and lower basal and length dependent %DRX and less calcium-stimulated thick filament activation shows these behaviors are critical to the decompensated RV. Precision targeting myotropes that impact these coupled behaviors may particularly improve those patients with these defects. This resonates with the recent clinical experience with omecamtiv mecarbil that had modest overall effects but greater impact in HFrEF patients with the lowest EF^41^ and worse functional capacity^42^. We suspect other methods to stimulate sarcomere function may achieve the desired results even more impactfully, and this work should help focus such efforts to improve HFrEF-PH therapy.

## ACKNOWLEDGEMENTS

This research used resources of the Advanced Photon Source, a U.S. Department of Energy (DOE) Office of Science User Facility operated for the DOE Office of Science by Argonne National Laboratory under Contract No. DE-AC02-06CH11357. This project was supported by grant P30 GM138395 from the National Institute of General Medical Sciences of the National Institutes of Health, R01 HL148785 (K.S.C), R01HL149891 (K.B.M), K23HL146889 (S.H.) and R35 HL 135827 (D.A.K.) from the National Heart Lung and Blood Institute of the National Institutes of Health, and support from the American Heart Association (V.J.). D.N. was supported by Department of Education GAANN Grant #P200A190080 to the Center for Synchrotron Radiation Research and Instrumentation (CSRRI) at the Illinois Institute of Technology. T.I provides consulting services to Edgewise Therapeutics Inc. and receives collaborative research funding from Bristol Myers Squibb Inc., but such work is unrelated to the content of this article.

