## Supplemental Materials for "Right Ventricular Sarcomere Contractile Depression and the Role of Thick Filament Activation in Human Heart Failure with Pulmonary Hypertension"

**Jani, et al.**

**SUPPLEMENTAL METHODS**

***Isometric tension-calcium, tension-length, and resting tension.***

Permeabilized (skinned) cardiomyocytes were prepared by cutting frozen tissue over dry ice in 5-10 mg pieces and immediately incubating in ice-cold (0°C) isolation buffer (5.55 mM Na2ATP, 7.11 M MgCl2, 2 mM EGTA, 108.01 mM KCl, 8.91 KOH, 10 mM Imidazol, 10 mM DTT) with 0.3% Triton X-100 in the presence of protease (Sigma-Aldrich, MO) and phosphatase inhibitors (PhosSTOP, Roche, Germany), as previously described ^15,16^. Tissue was then homogenized at 7500 rpm (OMNI Digital Programmable Homogenizer, Kennesaw, GA) for 6-8 seconds in 2-second pulses over ice with minimal frothing to generate a skinned cellular preparation after 20 minutes incubation at 4°C. After washing in isolation buffer with Triton-X100, cardiomyocytes were affixed to a force transducer-length controller (Aurora Scientific, Canada) using ultraviolet-activated adhesive (Norland, NJ). Cells were then transferred into room temperature (~22°C) relaxing buffer (5.95 mM Na_2_ATP, 6.41 mM MgCl_2_, 10 mM EGTA, 100 mM BES, 10 mM CrP, 50.25 mM Kprop, protease inhibitor (Sigma-Aldrich, St. Louis, MO), 1 mM DTT). Sarcomere length (SL) was measured by Fourier transformation of digital images (IPX-VGA210, Imperx, FL) and adjusted by micro-manipulators (Siskiyou, CA).

Isometric tension-calcium relations were assessed at SL=2.1 μm. Force developed at bathing solution Ca^2+^ ranging 0.0 to 46.8 μM (46.8 μM solution composition: 5.95 mM Na_2_ATP, 6.20 mM MgCl_2_, 10 mM Calcium-EGTA, 100 mM BES, 10 mM CrP, 29.98 mM Kprop, protease inhibitor (Sigma-Aldrich, St. Louis, MO), 1 mM DTT) was divided by myocyte cross sectional area (CSA=π/5⋅a^2^, a=short axis dimension) to calculate tension (mN/mm^2^). Tension versus -log[Ca^2+^] plots were fit to the three-element Hill equation: T = T_max_ [Ca^2+^]*^nh^*/(EC_50_*^nh^*+ [Ca^2+^]*^nh^*), yielding maximum isometric tension (T_max_, mN/mm^2^), calcium sensitivity (EC_50_, μM), and cooperativity (Hill coefficient, n_h_). Active length-dependent tension was assessed in 3.8 μM Ca^2+^ at 2.0-2.5 μm SL. Resting tension was assessed in Ca^2+^=0 buffer over a similar SL range.

***Cardiomyocyte Contractile Kinetics*.**

The rate constant of tension redevelopment after acute crossbridge disruption (k_tr_), hyperbolic tension-velocity dependence, and tension-power relatons^3,17,18^ were measured. K_tr_ was acquired at 1.7 and 3.8 μM [Ca^2+^], SL = 2.1 μm. After steady-state force was achieved, SL was reduced by 20% for 20 ms followed by re-stretch to the initial length using servocontrol. Force redevelopment data were fit to a mono-exponential (MATLAB, Mathworks 2020) to calculate k_tr._ Tension-velocity and tension-power relationships were obtained by starting at SL=2.1 μm and 3.8 μM [Ca^2+^], and then exposing cells to 5-10% stepwise lower reduction starting at T_max_ for 3.8 μM Ca^2+^). Velocity of cell shortening was measured over 75 ms immediately following the change in tension using linear regression to account for internal viscous load (primarily titin), and resulting exponential dependence between length and time.^3^ Force from the force transducer output was used for all measurements to correct for errors in servo-control. Data were then fit to the hyperbolic Hill equation (T+a)(V+b) = (T_3.8_+a)b to determine V_max_ and P=Tb((T_3.8_+a)/(T+a)-1) and peak power, Pwr_max_, using MATLAB (Mathworks, 2020).

***Cardiomyocyte Resting Viscoelasticity.***

Resting cardiomyocyte viscoelasticity was quantified using step force (creep) and step length (recoil) protocols under relaxing conditions. In the absence of calcium, myocytes can be considered standard solids with a single elastic element, determined from the force-SL curve, in parallel with an elastic and viscous element in series. For the creep test, a force clamp was applied at a load equivalent to a 14% SL change (2.1 to 2.4 μm), while for the recoil test, a 14% SL change (2.1 to 2.4 μm) was applied. Both were fit to mono-exponential to determine creep and recoil time constants (Mathworks, 2020).

***Small Angle X-Ray Diffraction*.**

Small angle X-ray diffraction patterns^7^ were acquired at the BioCAT beamline 18ID at the Advanced Photon Source of Argonne National Laboratory in Illinois (X-ray beam energy 12 keV, 0.1022 nm wavelength, incident flux 10^13^ photons per second).^19^ The X-ray beam was focused to 150x30 μm and 250x250 μm at the sample position. Flash-frozen tissue pieces (10-15 mg) were cut in ice-cold pCa 8 relaxing solution (2.25 mM Na2ATP, 3.56 mM MgCl2, 7 mM EGTA, 15 mM sodium phosphocreatine, 91.2 mM Potassium Propionate, 20 mM Imidazole, 0.165 mM CaCl_2_, and protease inhibitor cocktail pH 7.0) and skinned for 1 hour in relaxing solution containing 1% Triton X-100 and 15 mM BDM at room temperature. Myocyte strips were dissected to a length of ~4x0.2 mm and affixed to aluminum T-clips at both ends. Preparations were suspended between two hooks in a customized chamber with two Kapton windows in the X-ray path. The preparation was lengthened to SL 2.1 μm by monitoring light diffraction patterns from a helium-neon laser (633 nm). Radiation damage was minimized by moving the X-ray exposure region for each pattern. X-ray patterns were collected at 2.1 μm SL at pCa 8. For a subset of patients, myocyte strips were exposed to 2 µM mavacamten (Selleck Chemicals, Houston, TX) and 5.95 mM dATP-containing solutions (Millipore Sigma, Burlington, MA) in lieu of ATP for 15 minutes; patterns were collected pre- and post- drug exposure. A MarCCD 165 detector (Rayonix Inc., Evanston IL) with 1s exposure time was used for acquisition. Between two and four patterns were collected for each condition per patient.

Data were analyzed using the open-source MuscleX v1.5 software package, developed at BioCAT.^20^ Equatorial reflections measured using the “Equator” protocol in the software as described previously.^7^ Briefly, the intensity trace along the equation is summed across the image and integrated to a one-dimensional intensity reflection. The 1,0 and 1,1 reflections were modeled as Gaussian function superimposed on a smooth background. Lattice spacing was calculated by measuring the distance between the 1,0 reflection and the origin. For pattern visualization, “Quadrant Folding” in MuscleX v1.5 was used to visualize the equatorial and meridional reflections.

***Myosin ATP Turnover Kinetics.***

The percentage of SRX myosin (%SRX) was determined with a single nucleotide turnover assay^20-22^ applied to skinned single cardiomyocytes. Myocytes were set to 2.1 μm SL in relaxing buffer, washed in rigor buffer (relaxing buffer without ATP or CrP) for 1 minute, incubated for 1 minute in rigor buffer with the fluorescent ATP analog 25 μM 2’-/3’-O-(N’-Methylanthraniloyl) adenosine-5’-O-triphosphate (a.k.a. mant-ATP, Enzo Life Sciences, Axxora LLC, Framingham, NY), and moved to room temperature relaxing buffer. As mant-ATP (excitation 352-402 nm, emission 417-444 nm) was hydrolyzed, fluorescence was acquired with a photomultiplier tube (Horiba / PTI 814 Photomultiplier Detection System) continuously at 100 Hz for 1000 seconds using a Nikon Eclipse Ti2 inverted microscope.

The acquired fluorescence decay from the pulse chase protocol is bi-exponential: the initial rapid phase is driven by DRX myosin ATPase activity of DRX myosin while the slow phase is driven by the myosin ATPase rate of SRX myosin). Calculating the relative contribution of each phase can in turn determine the proportion of DRX versus SRX myosin in a cell. The raw fluorescence decay signal was fileted with a second-order Savitzky-Golay filter, normalized, and fit to I = 1 – P_1_(1-e^-t/T1^) – P_2_(1-e^-t/T2^). P_1_ and T_1_ measure the fraction and rate of myosin with fast ATP turnover, respectively, while P_2_ and T_2_ measure the fraction and rate of myosin with slow ATP turnover. The proportion of SRX myosin is then determined by 2⋅P_2_, while the percentage of DRX myosin is 1-(2⋅P_2_). Background noise was limited using the IonOptix Cell Frame Adapter (CFA, Westwood, MA). Background was subtracted by measuring average photomultiplier tube voltage output in the surrounding relaxing buffer at the end of the assay for each cardiomyocyte. A subset of myocytes was subjected to stretch (2.1 to 2.4 μm) and incubation with 2 μM mavacamten for 15 minutes in relaxing solution. Paired fluorescence decay measurements were taken before and after each of these perturbations. All analysis was performed in MATLAB (Mathworks, 2020).

**Transcriptomic Analysis**

Isolation of mRNA and sequencing were conducted by Genewiz (Next Generation Sequencing, RNA Sequencing, Azenta Life Science; 150 bp paired-end reads, 350 M reads per sample). Total RNA was isolated from flash-frozen non-failing (n=9) and HFrEF (n=21) RV myocardium. mRNA was enriched with poly-A selection. For sequence alignment, Hisat2 (version 2.2.0) indices^23^ were built using the human genome GRCh38 (FASTA file from ENSEMBL version 81) with gene set annotation in gtf format. SAMtools^24^ was used for SAM to BAM compression, and featureCounts was used to count reads to individual genes with accepted read quality.^25^ Differential expression analysis was performed with DESeq2 with FDR < 1x10^-6^. Principal component analysis was performed in Python (sklearn).

**SUPPLEMENTAL FIGURES AND TABLES**

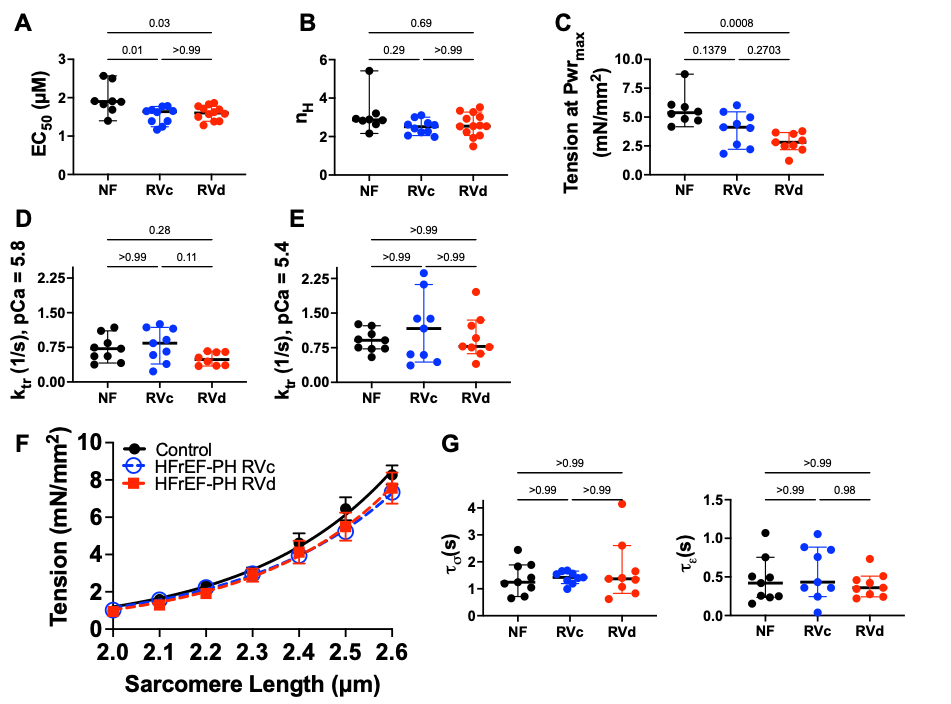

**Supplemental Figure 1. Calcium sensitivity, cooperativity, rate constant of tension redevelopment (k_tr_), resting tension, and viscoelastic time constants. (A)** Calcium concentration at 50% maximum tension (EC_50_) and **(B)** Hill Coefficient (n_H_) for control, RVc, and RVd HFrEF-PH subgroups identified by machine learning based on myocyte functional measures. P-values are from Kruskal-Wallis test with Dunn’s multiple comparisons. **(C)** Tension at maximum power determined from the derivative of the tension-power curve-fit using $P=\frac{\left( X_{0}+a \right)bT}{T+a}-bT$, where a,b, and X_0_ are fit parameters. X_0_ is equivalent to the isometric tension at Ca^2+^ = 3.8 µM. Tension at peak power is calculated from $\partial P/\partial T\left. \right|_{T=T_{opt}}=0$; yielding $T_{opt}=\sqrt{a\left( X_{0}+a \right)}-a$. P-values are from Kruskal-Wallis test with Dunn’s multiple comparisons. **(D, E)** Tension recovery time constant - K_tr_ for all three patient groups measured at 1.7 μM (half-activation) and 3.8 μM (maximum activation) Ca^2+^. P-values are from Kruskal Wallis test with Dunn’s multiple comparisons. (**F)** Resting tension for each group measured at 0 µM Ca^2+^ over sarcomere lengths ranging 2.0 to 2.6 μm. (**G)** Resting viscoelastic creep ($\tau_{\sigma}$) and recoil ($\tau_{\epsilon}$) for each patient group. Perturbations used to generate these time constants are rapid sarcomere length changes from 2.1 to 2.4 μm in the absence of Ca^2+^. P-values are from Kruskal Wallis test with Dunn’s multiple comparisons test.

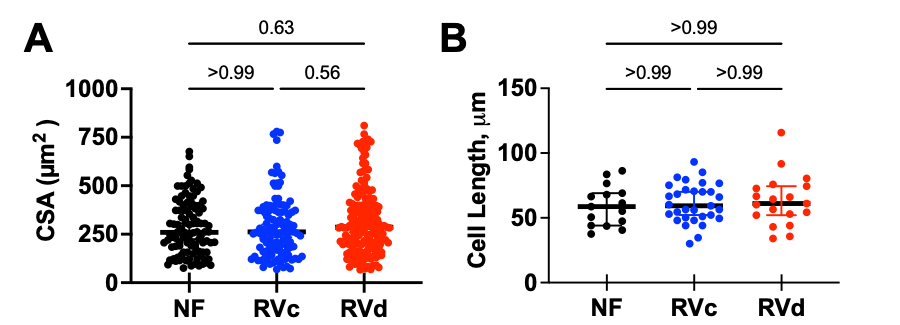

**Supplemental Figure 2. Cross sectional area and cell length for cardiomyocytes**. **A)** Cross sectional area (CSA) for all myocytes used for tension calculation calculations in this study for each patient group. Cross-sectional area is estimated as $\frac{\pi}{4}ab$, where a is the myocyte diameter directly measured by microscopic imaging, and b is the short axis diameter estimated as =0.8a. P values are from Dunn’s multiple corrections after a Kruskal Wallis test. **B)** Cell length of myocytes used for velocity-tension and power-tension measurements. This was directly measured from the microscopic imaging camera as the distance between the force and length transducer. Only myocyte measurements used for normalization are included. CSA was used for more replicates within all experiments (all isometric tension-ca, tension-length, drug incubations, and velocity/power measurements normalized measurements to CSA). Only velocity/power measurements depend on cell length, which included fewer technical replicates. P values are from Dunn’s multiple corrections after a Kruskal Wallis test.

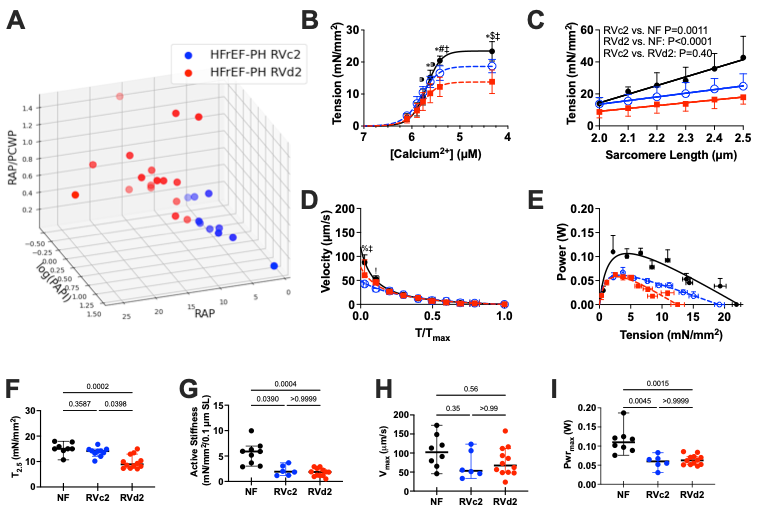

**Supplemental Figure 3. Machine learning trained on clinical features of RV functions predicts depressed isometric tension but not other measures of cardiomyocyte contractility.**

**A)** Three-dimensional scatter plot with right atrial pressure (RAP), right atrial to pulmonary capillary wedge pressure ratio (RAP/PCWP), and the logarithm of pulmonary artery pulsatility index (log(PAPi)), for each patient. Blue and red correspond to clusters derived by K-Means clustering of these clinical parameters – RVc2 (blue, RV compensation) and RVd2 (red, RV decompensation). Subsequent comparison of cardiomyocyte mechanical properties from these clinically determined subgroups are shown for **B)** tension-calcium, **C)** tension-length, **D)** tension-velocity, and **E)** tension-power relationships. RVd2 vs. Control: &P<0.05; ¶ P<0.01; %P<0.005; *P<0.0001; RVc2 vs. Control: !P<0.05; §P<0.01; #: P<0.005; $P<0.0001; RVd2 vs. RVc2: +P<0.05; ⁍P<0.01; ‡P<0.0001. RVc2- RV compensated (trained on clinical features). RVd2 – RV decompensated (trained on clinical features). Summary data for **F)** Isometric tension at 2.5 uM Ca^2+^, **G)** Active myocytes stiffness (length-tension slope), **H)** maximum shortening velocity (V_max_), and **I)** maximum power (Pwr_max_) quantified from the various relations shown in panels **B-E**.

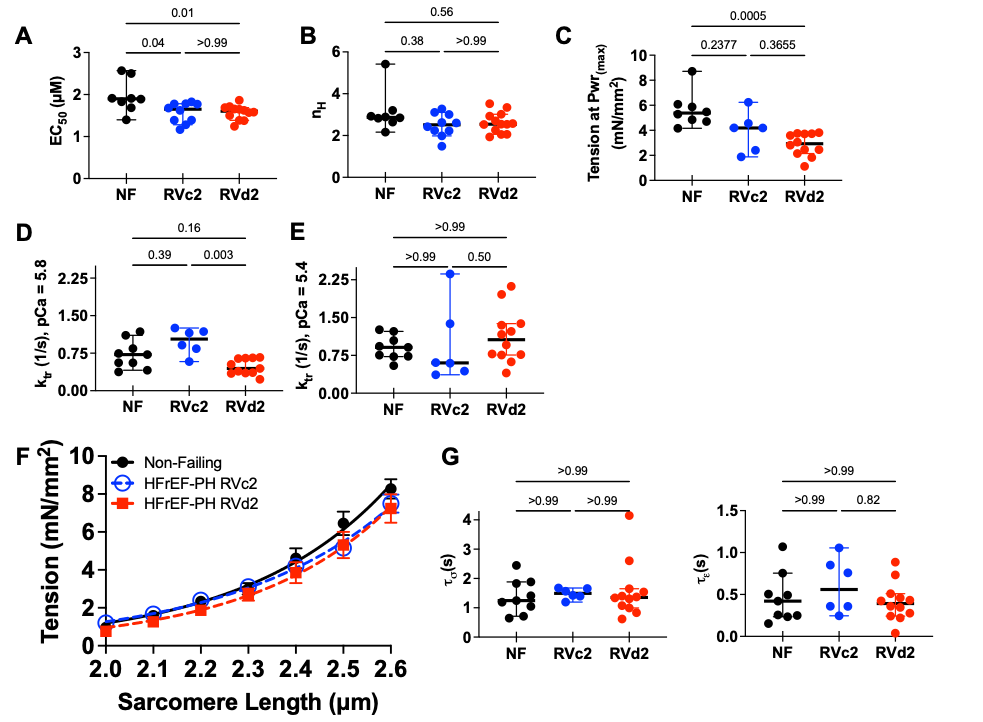

**Supplemental Figure 4. Calcium sensitivity, cooperativity, rate constant of tension redevelopment (k_tr_), resting tension, and viscoelastic time constants from ML based on clinical features. (A)** Calcium concentration for 50% maximum tension (EC_50_) and **(B)** Hill Coefficient (n_H_) for control, RVc, and RVd as identified by machine learning on myocyte functional measures. P-values are from Kruskal-Wallis test with Dunn’s multiple comparisons. **(C)** Tension corresponding to maximum power calculated from the tension-power fit. See analysis description in legend for Figure S1C. P values are from one-way ANOVA with Tukey multiple comparisons. **(D, E)** K_tr_ for three patient groups at 1.7 μM (half-activation) and 3.8 μM (maximum activation). P-values are shown from 2-way RMANOVA with Sidak’s multiple comparison between groups. (**F)** Resting tension for three patient groups calculated at 0 µM Ca^2+^ from 2.0 to 2.6 μm sarcomere length. (**G)** Resting viscoelastic creep ($\tau_{\sigma}$) and recoil ($\tau_{\epsilon}$) for control (black), RVc (blue), and RVd (red). Perturbations correspond to a sarcomere length change of 2.1 to 2.4 μm in the absence of Ca^2+^. P-values are from Dunn’s Multiple Comparison after a Kruskal Wallis test.

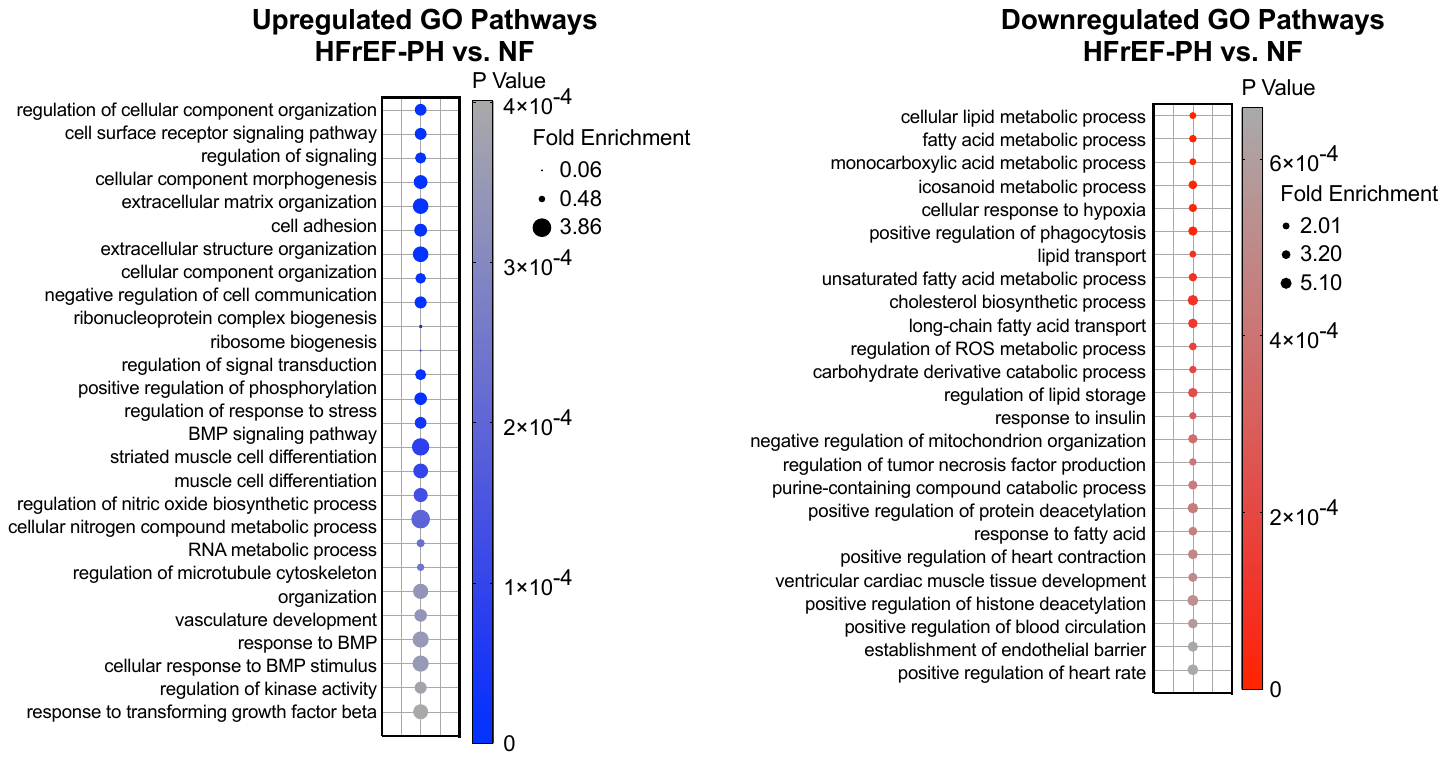

**Supplemental Figure 5. Pathway analysis for HFrEF-PH vs. control.** Gene ontology analysis on differentially expressed genes determined by RNAseq for HFrEF-PH myocardium versus control. GO terms were curated and redundant terms eliminated. The top 25 upregulated pathways and top 20 downregulated pathways are shown. Color corresponds to Bonferonni adjusted P-value and size corresponds to fold enrichment.

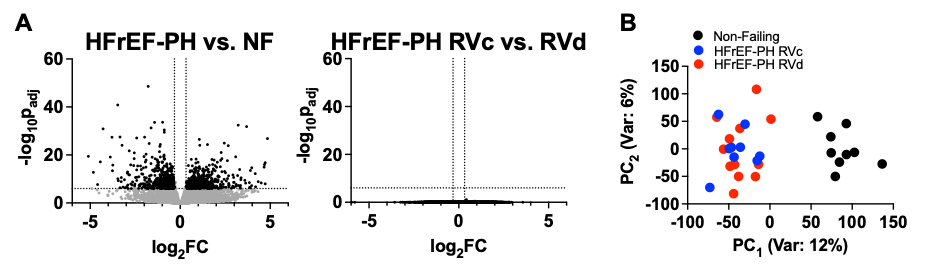

**Supplemental Figure 6. Transcriptomic and targeted sarcomere gene expression analysis in HFrEF-PH. A)** Volcano plots for ***left:*** HFrEF-PH vs. control and ***right:*** HFrEF-PH RVc vs. RVd. The latter reveals no significant expression changes between the two groups of HFrEF-PH patients. **B)** Principal component analysis shows overlap between HFrEF-PH RVc and RVd, but a clear separation from HFrEF-PH and non-failing controls.
**Supplemental Table 1. Detailed Laboratory, CPET, Right Heart Catheterization, and Echocardiographic Data for HFrEF-PH.** Data presented as mean ± standard deviation. BNP – Brain Natriuretic Peptide, VO_2_ – Maximum Oxygen Consumption, VE – Ventilatory Equivalent for CO_2_, VCO_2_ – Carbon Dioxide Production, LV – Left Ventricle, RV – Right Ventricle, TAPSE – Tricuspid Annular Planar Systolic Excursion, PASP – Pulmonary Artery Systolic Pressure, RVLSS – Right Ventricular Longitudinal Systolic Strain, IS – Interventricular Septal Strain.

| **Characteristic** | **HFrEF-PH (n=25)** |
| --- | --- |
| **Laboratory Values** | |
| Pro-BNP, ng/mL | 4892±5887 |
| Creatinine, mg/dL | 1.2±0.4 |
| estimated Glomerular Filtration Rate , mL/min | 64±21 |
| Total Bilirubin, mg/dL | 1.1±0.9 |
| **Cardiopulmonary Exercise Testing** | |
| 6 Minute Walk Distance, meters (n=5) | 323±87 |
| Peak VO_2_, mL/min/kg (n=10) | 14±4 |
| Respiratory Exchange Ratio (n=9) | 1.1±0.1 |
| VE/VCO_2_ slope (n=7) | 37±8 |
| **Right Heart Catheterization** | |
| Systolic Blood Pressure, mmHg | 97±11 |
| Mean Arterial Pressure, mmHg | 76±11 |
| Heart Rate, bpm | 78±22 |
| Right Atrial Pressure, mmHg | 12±6 |
| Pulmonary Artery Systolic Pressure, mmHg | 48±11 |
| mean Pulmonary Artery Pressure, mmHg | 30±8 |
| Pulmonary Artery Pulsatility index | 3.7±5.3 |
| Pulmonary Capillary Wedge Pressure, mmHg | 22±6 |
| Right Atrial Pressure/Pulmonary Capillary Wedge Pressure | 0.5±0.2 |
| Pulmonary Vascular Resistance, Wood Units | 2.4±1.4 |
| Cardiac Output, L/min | 4±2 |
| Cardiac Index, L/min/m^2^ | 2.1±0.7 |
| **Echocardiographic Characteristics** | |
| LV Ejection Fraction, % | 22±10 |
| LV Mass Index, g/m^2^ | 149±43 |
| LV End Diastolic Dimension, cm | 6.3±1.4 |
| Mitral E/A | 1.9±0.7 |
| e’, m/s | 5.1±2.2 |
| E/e’ | 20±13 |
| Tricuspid Regurgitant Maximum Velocity, m/s | 89±133 |
| Mitral Regurgitation (moderate-severe) | 88% |
| Basal RV End Diastolic Dimension, cm | 4.0±0.5 |
| Inferior Vena Cava Diameter, cm | 2.5±0.7 |
| Tricuspid Annular Planar Systolic Excursion, cm | 1.6±0.5 |
| Right Atrial Area, cm^2^ | 76±104 |
| Right Atrial Volume index, mL/m^2^ | 80±44 |
| Fractional area change, % | 25±4 |
| Tissue Doppler Imaging S’, m/s | 9±4 |
| RV Wall Thickness | 0.8±0.1 |
| Echo RV Systolic Pressure, mmHg | 45±19 |
| TAPSE/PASP Ratio | 0.4±0.2 |
| Global RVLSS, % | -12±7 |
| Global RVLSS + IS , % | -9±5 |
